# PPM1D truncation-associated overexpression of the stress-related protein NQO1 confers sensitivity to the bioactivatable drug IB-DNQ in diffuse intrinsic pontine glioma

**DOI:** 10.1101/2024.09.05.611476

**Authors:** Maxime Janin

## Abstract

Diffuse intrinsic pontine glioma (DIPG) is a very aggressive brainstem tumor with poor survival and a lack of effective treatments. In this study, I observed the differential overexpression of the stress-related protein NAD(P)H Quinone Dehydrogenase 1 (NQO1) in some patient-derived DIPG cell lines and tumors. I sought to understand how this protein is regulated in DIPG and to investigate the therapeutic potential of the NQO1-bioactivatable drug Isobutyl-deoxynyboquinone (IB-DNQ). Interestingly, the study of the mutational profiles of the cell lines indicated that truncation of PPM1D correlated with NQO1 overexpression. From a functional standpoint, cellular models were utilized to unravel the link between PPM1D phosphatase and NQO1 expression in DIPG by dephosphorylating MDM2 serine 395, leading to NQO1 protein stabilization. From a therapeutic perspective, IB-DNQ treatment showed an NQO1-dependent growth inhibition sensitivity in vitro and induced an extended survival in vivo. Overall, my results reveal a new regulation of NQO1 at the protein level in PPM1D-mutated DIPG indicating a promising therapeutic approach.

## Introduction

Patients with paediatric Diffuse Intrinsic Pontine Glioma (DIPG) have very poor survival, and no effective treatment for the disease is available. DIPG is the main paediatric brainstem neoplasm, accounting for approximately 80% of brainstem tumours. It is mostly diagnosed in children between 5 and 10 years of age, and more than 90% of patients die within 2 years from diagnosis [34,38]. DIPG originates in the pons, a critical part of the brainstem, and diffusely infiltrates this very sensitive region controlling vital functions such as the regulation of respiration or involuntary functions, making its complete resection by surgery not feasible. Recurrent mutations in the genes encoding the H3.3 (gene *H3F3A*) and H3.1 (genes *HIST1H3B* and *HIST1H3C*) histone variants, resulting in lysine-to-methionine substitution at position 27 (K27M), occur in more than 85% of DIPGs [17,28]. Other mutations have also been described as main drivers of DIPG development such as those present in *PPM1D* (protein phosphatase, Mg^2+^/Mn^2+^ dependent 1D)*, ACVR1* (activin A receptor type 1)*, PIK3R1* (phosphoinositide-3-kinase regulatory subunit 1) and *p53* [17,28]. The unique biology and clinical features of DIPG make therapeutic approaches for other paediatric brain cancer ineffective. Despite numerous clinical trials, the median overall survival remains around 12 months after diagnosis, with radiotherapy providing a 3 month survival benefit [10]. Therefore, it is urgent to increase our understanding of the biology of DIPG and identify new treatment strategies to tackle this paediatric cancer.

It is known that tumors, including gliomas, use different pathways to mitigate oxidative stress-associated cell death. In this study, I describe that in patient-derived DIPG cell lines and biopsies PPM1D-truncation mediates dephosphorylation of MDM2 that upregulates the multifactorial stress protein NQO1. Most important, I observed that DIPGs overexpressing NQO1 were very sensitive to the antiproliferative effect of the NQO1-bioactivatable substrate IB-DNQ, a drug that crosses the blood-brain barrier [15] and it is well-tolerated in preclinical models [25,26]. Thus, I propose to assess the effect of IB-DNQ in preliminary clinical assays as a candidate therapeutic approach against PPM1D-truncated DIPG. This strategy could prove beneficial for DIPG patients.

## Materials and methods

### Cell lines

Human DIPG patient’ derived cell lines HSJD-DIPG-007; HSJD-DIPG-008; HSJD-DIPG-011; HSJD-DIPG-012; HSJD-DIPG-013 and HSJD-DIPG-014 were obtained from the Sant Joan de Deu paediatric hospital in Barcelona (Catalonia, Spain); SU-DIPG-VI was obtained from Dr. Michelle Monje’s laboratory in Stanford (Stanford Medicine; USA). Cell lines were all cultivated in a 1:1 mixed Neurobasal-A medium with DMEM/F12 medium supplemented with HEPES buffer, sodium pyruvate MEM, Non-essential amino-acids, GlutaMax-I, antibiotic-antimycotic, B-27 supplement without vitamin A (all from ThermoFisher), human PDGF-AA, human PDGF-BB, recombinant human FGF, recombinant human EGF (all from Peprotech) and heparin solution 0,2% (Sigma). All cell lines were tested for the absence of mycoplasma.

### Expression Analyses

For real time quantitative reverse transcription PCR experiments, total RNA was extracted using the SimplyRNA kit (Promega) on a Maxwell RSC device (Promega) and retrotranscribed using the ThermoScript RT-PCR system (ThermoFisher) with oligo(dT) primers. The reaction was carried out following the methods for use of SYBR Green (Applied Biosystems), and GAPDH was used as housekeeping gene to enable normalization. Primers for RT-PCR are listed in **Key Resources Table**. For immunoblotting assays, I extracted cell pellets and brain white matter samples with Laemli Buffer, sonication and 5 minutes at 95°C. For immunohistochemical analysis, paraffin-embedded sections of DIPG tumors were immunostained with the NQO1 antibody. Antibodies used in this study are described in **Key Resources Table**.

### Drug assays

IC50 studies were performed using the 3-(4,5-Dimethyl-2-thiazolyl)-2,5-diphenyl-2H-tetrazolium bromide (MTT) assay in drug-treated DIPG cell lines. Briefly, 48h after exposure to nine increasing concentrations of the drug, 10uL of diluted MTT was added to culture medium and incubated in culture incubator for 4 hours. Then, 100uL of Lysis buffer was added to each well and the plates were incubated overnight in the incubator. The days after, the 540 nm-optical densities were determined using a microplate reader (Perkin Elmer Viktor 3). Drugs used in this study are listed in **Key Resources Table.**

### DNA methylation analyses

The DNA methylation array used was the Infinium MethylationEPIC BeadChip (Illumina). Raw fluorescence intensity values were normalized with Illumina Genome Studio software (V2011.1) using ‘control normalization’ with background correction. Normalized intensities were then used to calculate DNA methylation levels (beta values).

### Gene transfection

The cDNA sequences of NQO1, PPM1D or MDM2 were cloned either into the pLVX-IRES-ZsGreen1 or pLVX-IRES-TOMATO expression plasmid (Clontech Laboratories) between defined restriction sites. Lentiviruses containing this construct were produced by cotransfecting HEK-293T cells with the recombinant plasmid, with packaging vectors and jetPRIME® Transfection Reagent (Polyplus Transfection) following the supplier instructions. Briefly, 10ug of each encoding plasmid was mixed with 7.5μg of ps-PAX2 and 2.5μg of PMD2.G plasmid (Addgene). Upon 10 min of RT incubation, the transfection mix was added dropwise on a 10 cm culture plate containing HEK293-TLV lentiviral packaging cells at 80% confluence. The transfection cocktail was removed after 6h and replaced by fresh medium. After 72h, viral-containing supernatant was collected, 0.45µM-filtred and stored at 4°C before infection. The recombinant product was randomly inserted by lentiviral transduction in the genome of the DIPG cell lines. After 5 passages, the green or red fluorescent cells were sorted by FACS and cultured in the above-described DIPG-specific medium. For primers sequences please refer to **Key Resources Table**.

### Gene depletion

Four different sequence gene-specific short hairpin RNA molecules (shRNAs) for NQO1 mRNA were designed and transduced into DIPG cell lines (for sequences please refer to **Key Resources Table**). shRNA against the MSS2 yeast mRNA (not present in mammals) was used as a scramble control. All shRNA molecules were ligated into pLVX-shRNA2-ZsGreen plasmid (Clontech). Lentiviral particles were produced following the same steps than in the previous section. Infected cells were FACS-sorted after 5 passages and cultured in DIPG-specific medium.

### Brain tumor xenografts

For tumor growth into the brainstem pons area, 1 x10^6^ of HSJD-DIPG-007 cells were inoculated into the pons of 10-9 week-old athymic Nude-Foxn1nu mice (Charles River Laboratories) for each treatment condition. All mouse experiments were approved by and performed according to the guidelines of the Xenopat company in agreement with the European Union and the Spanish national directives. The experiments were also approved by the local ethics committee. Globally, the “Principles of laboratory animal care” (NIH publication No. 86-23, revised 1985) were followed.

After 20 days, half of the mice were randomly chosen to be treated with IB-DNQ intravenously at 12mg/kg every 2 days for 10 days (5 doses) while the other half were treated with vehicle (hydroxypropyl beta-cyclodextrin (HPbCD)).

### Patients’ samples

Tumor samples were obtained from autopsies, freshly frozen, and collected at Sant Joan de Deu paediatric hospital, Stanford Monje Lab, and Washington D.C. paediatric hospital. A specific Material Transfer Agreement (MTA) was established between the different institutions and the Josep Carreras Institute, and the institutions declared written consent established with the family of the donor.

### Data availability

Main data generated for this study are presented in the two datasets.

## Results

### Overexpression of the NQO1 protein in patient-derived DIPG cell lines

Oxidative stress-induced death of cancer cells is a barrier to tumor growth and transformed cells have devised different ways to bypass this shortcoming [27]. One mechanism involves the induction of proteins that regulate the production and oxidative stresses. In this regard, NQO1 is a relevant multifunctional protective antioxidant that confers great protection to cellular death [33]. Overexpression of the stress-related protein NQO1 has been described in several cancer types, including pancreatic, lung, breast, thyroid, adrenal, ovarian, colon and corneal tumors [11,16,35,36]. Its high expression is usually associated with poor prognosis [4,8,22]. The molecular pathways leading to NQO1 overexpression in human malignancies are not completely understood and so far, no study of NQO1 protein expression have been performed in paediatric brainstem tumors. Using as starting biological material six patient-derived DIPG cell lines (all of them H3K27M-mutated), I observed by western-blot high expression of NQO1 in three cases (**Fig. 1a**). The remaining three DIPG cell lines (**Fig. 1a**) and control human astrocytes and brain white matter samples (**Suppl. Fig. S1a**) showed low levels of the NQO1 protein. The different amount of NQO1 was not associated to distinct DNA methylation patterns in the NQO1 promoter CpG island (**Suppl. Fig. S1b**) or in the NQO1 regulator NSUN5 (**Suppl. Fig. S1c**) [15]. Importantly, this was not just an *in vitro* phenomenon, since the use of primary autopsy DIPG samples confirmed that NQO1 overexpression determined by immunohistochemistry was also detectable. One block, N1327, corresponded to the autopsy that led to the derivation of the HSJD-DIPG-008 cell line, and this tumor had one of the highest expressions (**Fig. 1b**). Interestingly, the NQO1 RNA expression levels among the six DIPG cell lines were not statistically different (**Fig. 1c**), suggesting that the translocation of the transcription factor NRF2 into the nucleus [21], which leads to an increase of RNA transcription of different stress-related genes, including NQO1, was not involved either. Thus, protein translational or posttranslational regulation could be involved to generate the different NQO1 protein levels observed. To know if NQO1 protein upregulation was due to different protein stability, I used the cycloheximide chase analysis of protein degradation in the studied DIPG cell lines. Western blot analysis of NQO1 level did not show loss of protein stability upon cycloheximide treatment (**Fig. 1d**), suggesting that distinct protein levels could be associated to different translation rates. Indeed, variation on NQO1 protein levels upon cycloheximide treatment was similar in all cell lines, displaying a time-dependent increase on NQO1 levels, which suggests distinct translation rates instead of differential protein degradation.

**Figure 1.**
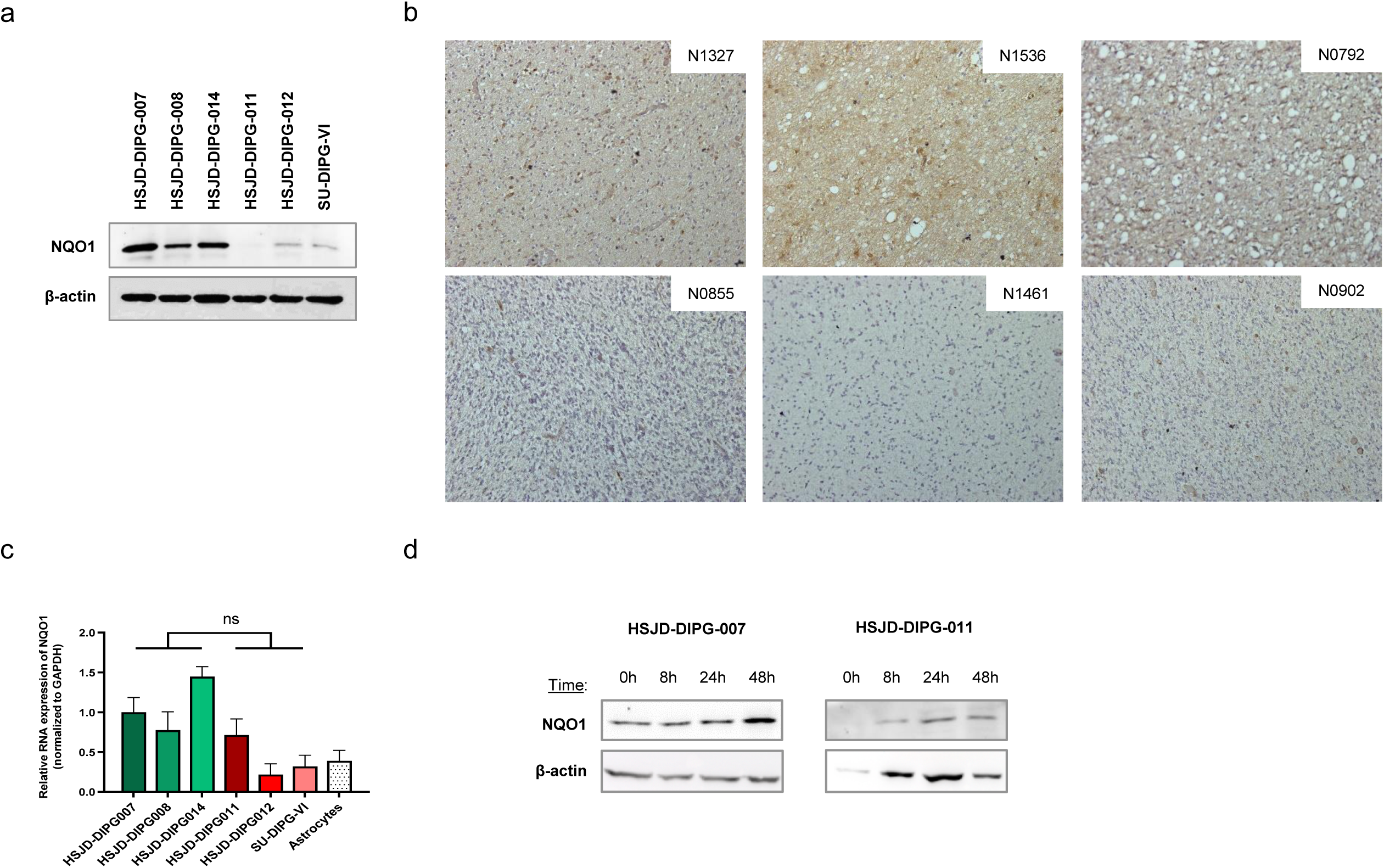
Expression of NQO1 in DIPG cell lines. (**a**) Western blot of NQO1 in the 6 DIPG cell lines. (**b**) Representative images of immunodetection of NQO1 in 6 Formalin-fixed paraffin embedded sections with same optical zoom and light exposure. Sections with highest expression of NQO1 are up and sections with lowest expression are down. Section corresponding to HSJD-DIPG-008 cell line is N1327 up on the left (**c**) NQO1 expression levels in DIPG cell lines and astrocytes control determined by real-time PCR (data shown represent mean ±s.d. of biological triplicates). (**d**) Western blot of NQO1 and beta-actin after cycloheximide treatment at 500 uM during different times in HSJD-DIPG-007; -008; -011 and -012. For real time PCR (c) significance was calculated using an unpaired t-test. ns: not significant. All drug-response curves, real time PCR and relative expression bar plots were generated using GraphPad Prism software in this study.

### PPM1D is mutated in DIPG cell lines with high expression of NQO1

To address which molecular settings could induce an enhanced translation rate leading to NQO1 protein overexpression, I wondered about the genetic context of the six studied DIPG cell lines beyond its shared H3K27M-mutated status. Remarkably, the 3 DIPG cell lines with high NQO1 protein expression levels were mutated for the PPM1D gene (also known as WIP1) and the 3 others with low NQO1 protein expression levels were wild type (**Fig. 2a**) [5]. Moreover, the expression of NQO1 in PPM1D mutated freshly frozen autopsy tumours was higher than wild-type DIPG cases or normal cortex (**Fig. 2b**). Further supporting the presence of NQO1 overexpression in the primary tumour of origin. The normal cortex was a matched sample from the patient with the SU-DIPG-35 PPM1D tumour, confirming the cancer-specific event of the NQO1 upregulation.

**Figure 2.**
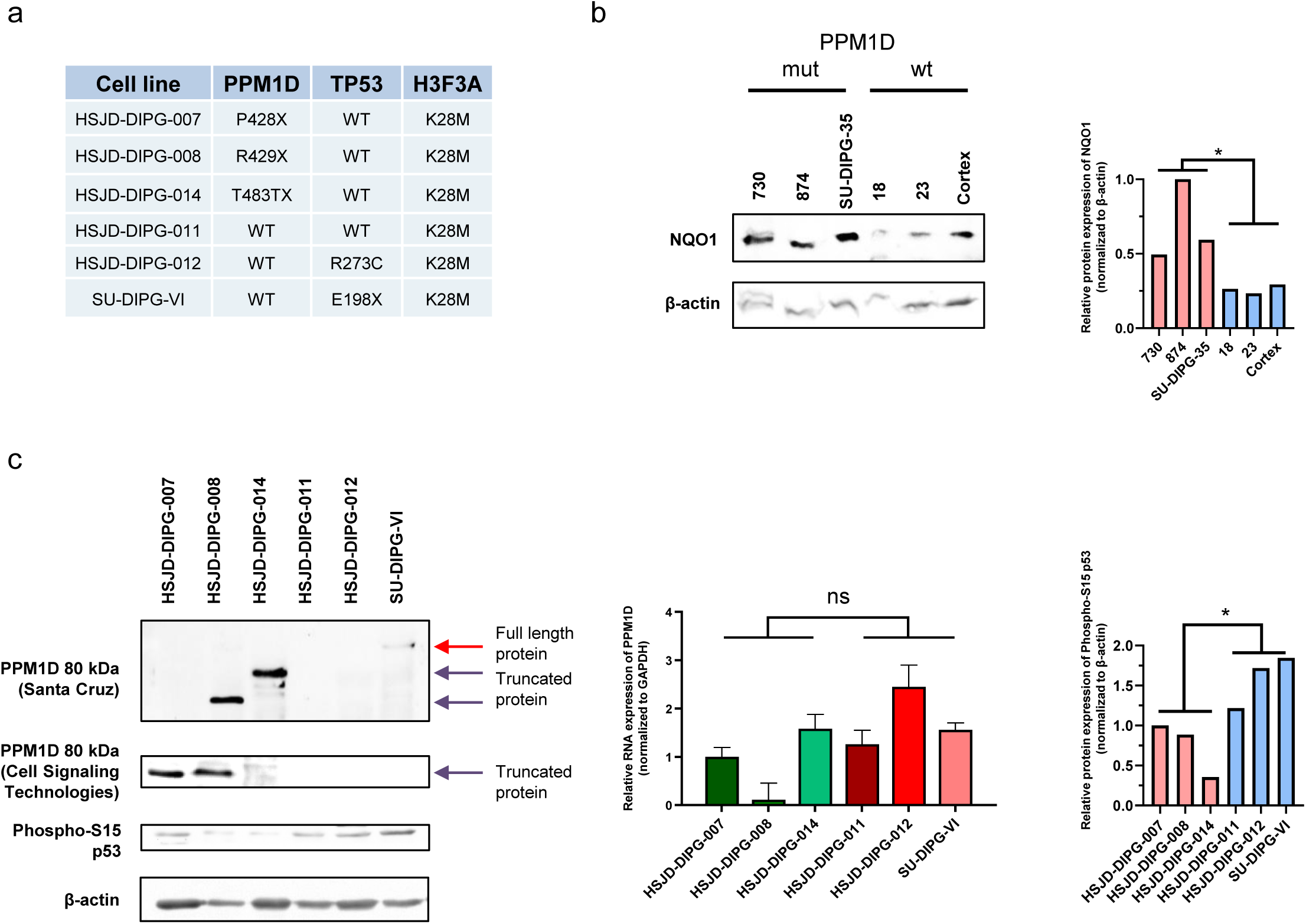
Patient-derived cell lines HSJD-DIPG-007; HSJD-DIPG-008 and HSJD-DIPG-014 are mutated for PPM1D. (**a**) Mutational status of the 6 DIPG cell lines. (**b**) Western blots of NQO1 in tumors from patients with or without PPM1D mutation. ( **c**) Left: Western blot of PPM1D showing accumulation in the 3 DIPG cell lines that overexpress NQO1 and western blot of P53 (S15) showing a slight variation between the two groups of cells. Middle: Real time PCR of PPM1D in the 6 DIPG cell lines studied. Right: Intensities of phospho-S15 p53 relative to beta-actin. For (b, c, and d) significance was calculated using an unpaired t-test. ns: not significant and * P<0.05.

PPM1D is mutated in approximately 20% of DIPG tumours [42] with point mutations in the sixth exon that result in protein truncation, leading to the production of C-terminal domain-lacking proteins. The loss of this domain causes a protein accumulation that is still functionally active inside the cells [18,20], and impairs DNA damage response [9]. As expected, I observed by western blot, using different antibodies depending on epitope recognition, the accumulation of PPM1D in the three mutant DIPG cell lines (**Fig. 2c**). PPM1D RNA levels did not show differences among the cell lines (**Fig. 2c**). Interestingly PPM1D dephosphorylates and destabilizes p53 [24]; whereas NQO1 stabilizes p53 [2,32]. Thus, PPM1D and NQO1 share a common target but with opposite activities and, thus, I wondered if the observed NQO1 overexpression was an attempt of the cells to compensate against PPM1D accumulation and over activity. In this regard, I observed a decrease of PPM1D mediated p53 S15 phosphorylation in those DIPG cells overexpressing NQO1 (**Fig. 2c**).

### Mutant PPM1D regulates NQO1 expression in DIPG cell lines

To assess if PPM1D truncation had a direct effect on NQO1 expression, I constructed different DIPG genetically engineered cellular models ( **Suppl. Fig. S2**). I observed that the stable transfection of the SU-DIPG-VI cell line with the PPM1D truncated form induced an increase in NQO1 protein levels (**Fig. 3a**). NQO1 upregulation was not observed upon transfection of the full-length form ( **Fig. 3a**), a mutant inert form of the full-length (**Suppl. Fig. S2a**) or a mutant inert form of the truncated PPM1D protein (**Suppl. Fig. S2a**). Conversely, I also studied the loss of the truncated PPMD1 protein on NQO1 expression by generating a PPM1D knock-out cell model (**Suppl. Fig. S2b**) by CRISPR-Cas9 in the PPM1D truncated mutant HSJD-DIPG-007 DIPG cells (**Fig. 3b**). I observed that, upon efficient abrogation of PPMD1 expression, there was a reduction of NQO1 protein levels (**Fig. 3b**). Interestingly, PPM1D inhibition by small molecules have been suggested as a possible treatment in DIPG to reactivate DNA damage response and decrease tumor growth [1]. Thus, I assessed if the use of the PPM1D inhibitor GSK2830371 in the PPM1D truncated mutant HSJD-DIPG-007 and HSJD-DIPG-014 DIPG cell lines could lead to NQO1 decreased expression and this was indeed the case (**Fig. 3c**). I also confirmed the regulation of the truncated form of PPM1D on NQO1 protein expression in a recently developed genetically modified mouse model [19]. I observed that endogenous truncation of *Ppm1d* ^exon^ ^6^ in brainstem tumors was associated with an increase in NQO1 protein levels (**Fig. 3d**), but not in the corresponding RNA transcript (**Fig. 3d**). In addition, the analysis of mouse neural stem cell (mNSC) transfected with control GFP or PPM1D-truncated plasmids demonstrated NQO1 overexpression at the protein level in PPM1D transfected cells, but not for the RNA transcript (**Fig. 3e**).

**Figure 3.**
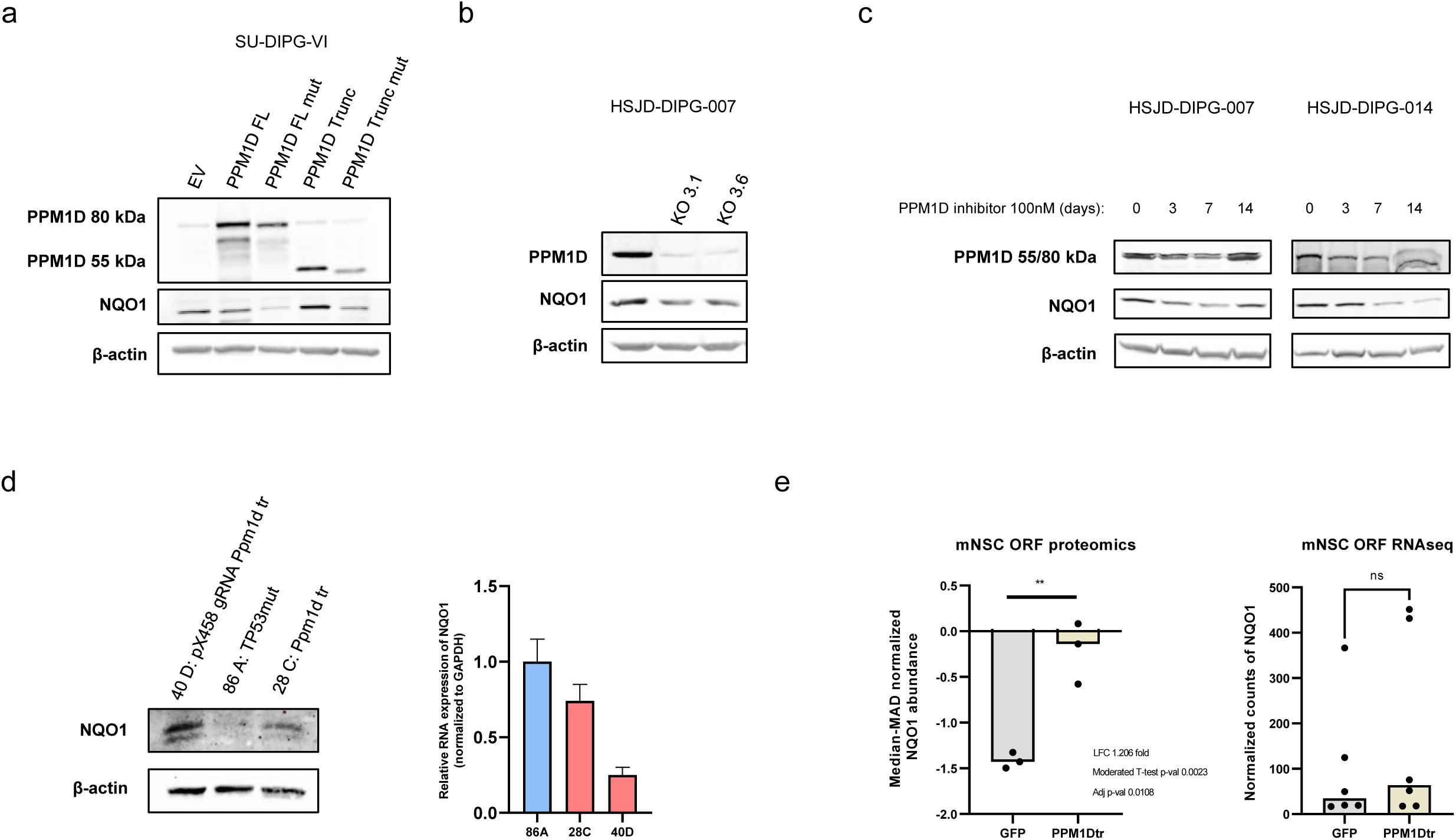
NQO1 expression is modulated by PPM1D. (**a**) Western blot analyses of PPM1D overexpression in SU-DIPG-VI cell line 6 weeks after stable transfection. EV: empty vector; FL: full length; mut: mutated; Trunc: truncated. (**b**) Western blot of PPM1D and NQO1 in HSJD-DIPG-007 KO cell lines. (**c**) Western blot of PPM1D and NQO1 in HSJD-DIPG-007 and HSJD-DIPG-014 cell lines treated at 100nm during 0; 3; 7 or 14 days. (**d**) NQO1 expression in mouse cell lines derived from IUE tumors analysed by western-blot (left) and real time PCR (right). (**e**) Proteomic and RNA-seq expression of NQO1 in mouse Neural Stem Cells (mNSC) transfected with GFP as control or with PPM1D truncated. For (e) significance was calculated using a moderated t-test. ns: not significant and ** P<0.005.

### Dephosphorylation of MDM2 at serine 395 by truncated PPM1D leads to NQO1 protein upregulation

Overall, the above-described data indicated that the truncated form of PPM1D is a key regulator of NQO1 protein levels in DIPG, but at that stage a direct effect or a phosphatase activity-mediated effect of PPM1D on an intermediate downstream target could not be discerned. One of the most well-known targets of PPM1D is the oncoprotein MDM2 (Mouse double minute 2 homolog) [23], a negative regulator of p53 protein levels. In this regard, the mutations in the phosphatase PPM1D leads to dephosphorylation of MDM2 at the S395 residue; a change associated with MDM2 stabilization and enhanced interaction-induced degradation of p53 [23]. I wondered if the PPM1D dependent-phosphorylation status of MDM2 could have additional roles beyond p53, for example acting through NQO1. Thus, I constructed a serine 395 to alanine mutant (S395A) MDM2 mutant plasmid for stable transfection in DIPG PPM1D wild-type cells. I observed that the transfection of this S395A mutant, that mimics the PPM1D phosphatase activity, in HSJD-DIPG-011 cells increased NQO1 protein expression level (**Fig. 4a**) but did not induce changes of NQO1 mRNA expression (**Fig. 4a**). Thus, to avoid excessive degradation of p53 and cell death in DIPGs, it is possible that PPM1D-mediated dephosphorylation of MDM2 leads to the upregulation of NQO1, a p53 stabilizer [2]. In this regard, MDM2 regulates protein translation [13,29] and this mechanism could be invoked herein, since changes in NQO1 protein degradation were not observed (**Fig. 1d**).

**Figure 4.**
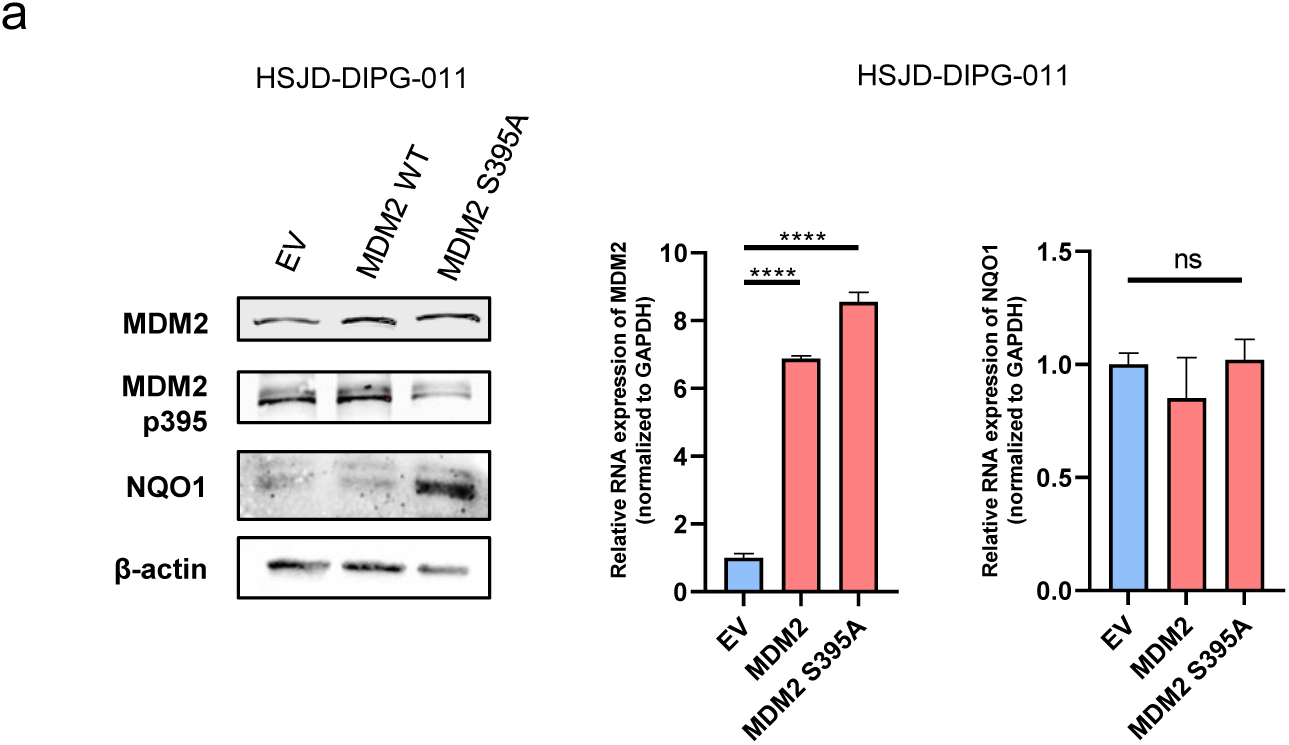
MDM2 serine dephosphorylation leads to NQO1 protein increased expression. (**a**) Stable transfection in HSJD-DIPG-011 of MDM2 S395A validated by western blot leads to NQO1 increased expression analysed by western blot (left), but do not modulate the transcript expression of NQO1 analysed by real time PCR (right). For real time PCR, significance was calculated using an unpaired t-test. ns: not significant and ****P<0.0001.

### NQO1 overexpression in DIPGs with PPM1D truncation-mutations are sensitive to the bioactivatable drug IB-DNQ

The reduction of quinone substrates by NQO1 to their respective hydroquinones in a futile redox cycle results in a high production of reactive oxygen species (ROS), causing extensive DNA lesions, PARP1 hyperactivation, and severe NAD+/ATP depletion that stimulate Ca2+–dependent programmed necrosis [14]. This mechanism has proved to be useful for inducing an outsized oxidative stress level that leads cancer cells to their death [31]. In this regard, various drugs have been found to be NQO1-bioactivatable, such as Beta-lapachone (ARQ 501) or quinone derivates as deoxynyboquinone and isobutyl-deoxynyboquinone (DNQ and IB-DNQ, respectively), which have demonstrated their anticancer activity in cancer cell line models [3,31,40]. Importantly, the compound IB-DNQ has demonstrated excellent pharmacokinetics data and a safe tolerability in domestic feline species models that further support the preclinical development of this drug [25,26]. Thus, I wondered if those DIPGs harboring PPM1D truncation-associated NQO1 overexpression showed a particular sensitivity to IB-DNQ.

Using the MTT assay to determine the IC50 cell growth inhibition values in my panel of DIPG cell lines, I observed that the 3 DIPG cell lines with PPM1D truncating mutation and associated high NQO1 expression (HSJD-DIPG-007; -008 and -014) were significantly more sensitive to the drug than the 3 DIPG cell lines with wild-type PPM1D and low levels of NQO1 (HSJD-DIPG-011; HSJD-DIPG-012 and SU-DIPG-(z test, P < 0.0001) (**Fig. 5a**). The use of different NQO1 constructs in these cells further confirmed this link. The development of stable depletion models of NQO1 by transfection of short hairpin RNA (shRNA) in NQO1 high-expressor HSJD-DIPG-007 cells showed an acquired resistance to the growth inhibition effect mediated by IB-DNQ (**Fig. 5b**). On the opposite side, the induction of NQO1 overexpression by stable transfection in cells with originally low-levels of NQO1 (HSJD-DIPG-011) rendered these cells to a more sensitive growth inhibitory effect upon IB-DNQ use (**Fig. 5c**).

**Figure 5.**
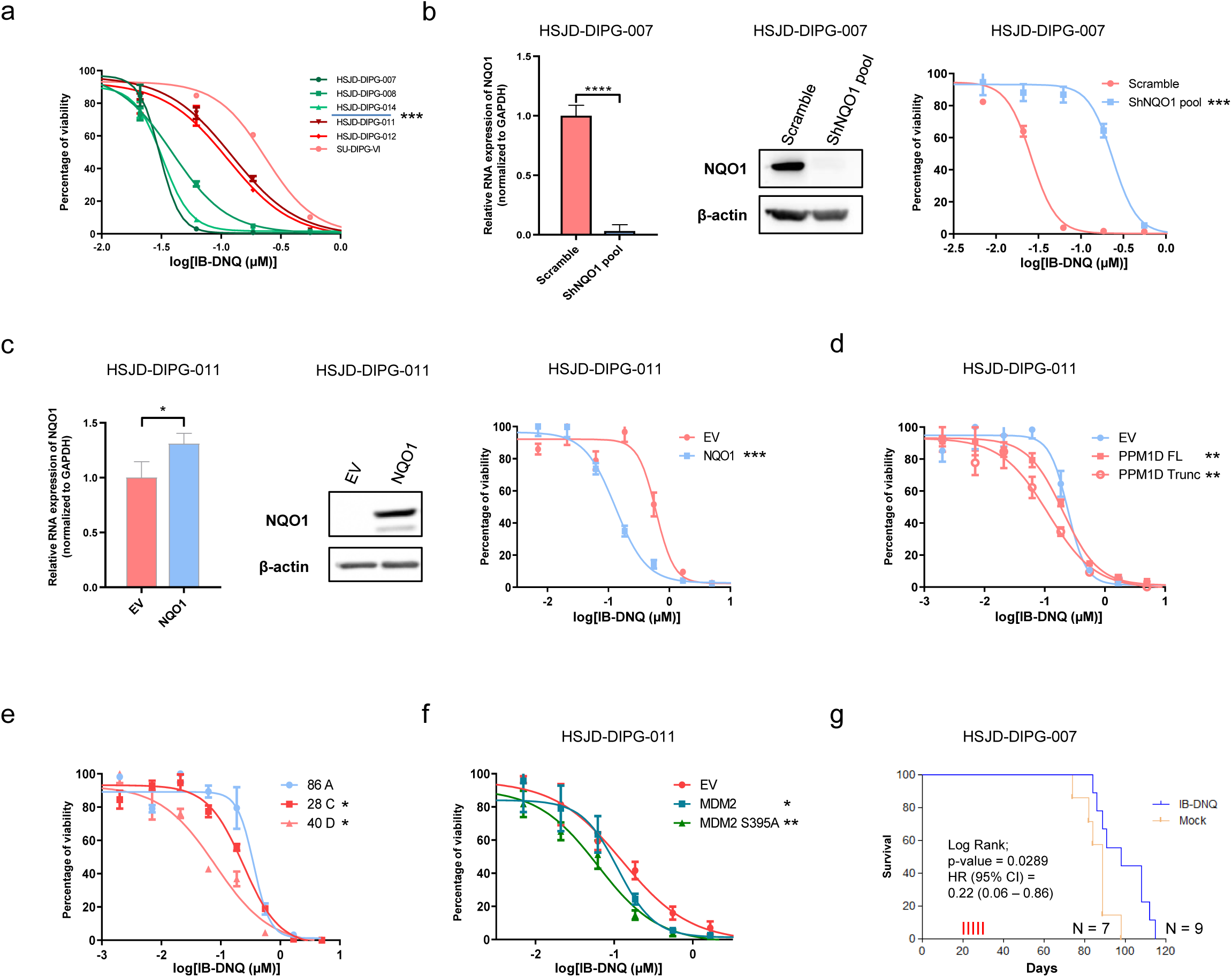
NQO1 modulation leads to IB-DNQ sensitivity in DIPG. (**a**) IC50 determination using the MTT assay in the DIPG cell lines. DIPG cells harbouring NQO1 overexpression (HSJD-DIPG-007; -008 and -014) show higher sensitivity to IB-DNQ in comparison to NQO1 normal expression cells (HSJD-DIPG-011; -012 and SU-DIPG-VI). (**b**) DIPG cellular models of NQO1 depletion by short hairpin (sh) RNA validated by real time PCR (left panel) and western blots (middle) for the cell line HSJD-DIPG-007. NQO1 depletion leads to IB-DNQ resistance shown by IC50 determination (right panel). (**c**) NQO1 overexpression model by stable transfection in HSJD-DIPG-011 cell line validated by real time PCR (left panel) and western-blots (middle) that leads to IB-DNQ sensitivity (right panel). (**d**) PPM1D overexpression by stable transfection in HSJD-DIPG-011 cell line leads to IB-DNQ sensitivity. (**e**) Cell lines derived from IUE mouse models of PPM1D are more sensitive to IB-DNQ. (**f**) MDM2 and mainly MDM2 S395A stable transfection in HSJD-DIPG-011 lead to IB-DNQ sensitivity. (**g**) Kaplan-Meier analysis of survival according treatment conditions (IB-DNQ treated vs mock group) in a mouse model with implanted tumors derived from the NQO1 overexpressing DIPG cell line DIPG 7. Mice was treated at days 20; 22; 24; 26 and 28 (red vertical lines). Significance of the log-rank test is shown. Results of the univariate Cox regression analysis are represented by the Hazards Ratio (HR) and 95% Confidence Interval (CI). Significance was calculated using a paired t test for IC50s and an unpaired t-test for real time PCR. *P<0.05, **P<0.005, ***P<0.0005 and ****P<0.0001.

The involvement of the PPMD1 truncation mutation leading to NQO1 upregulation and, thus, being itself a biomarker of response to the IB-DNQ compound was also demonstrated for the herein described genetically engineered human and mice models. In this regard, I observed that the stable transfection of the PPM1D truncated form in non-mutated DIPG cell lines not only induced an increase in NQO1 protein levels (**Fig. 3a**), but also increased the sensitivity to IB-DNQ (**Fig. 5d**). The use of the mouse model with a truncated form of PPM1D [19], where I have shown above the induced NQO1 overexpression (**Fig. 3d**), further confirmed that the developed brainstem tumors were highly sensitive to IB-DNQ in comparison to non-truncated PPM1D specimens (**Fig. 5e**). These observations were extended to the target of the PPM1D truncation, the oncoprotein MDM2. I found that transfection of the S395A MDM2 mutant (that mirrors he PPM1D phosphatase activity) in DIPG PPM1D wild-type cells not only induced NQO1 upregulation (**Fig. 4a**), but also enhanced the sensitivity to the IB-DNQ drug (**Fig. 5f**). As DIPG originates into the pons, a sensitive area into the brain that render difficult therapeutics’ delivery [39], I tested the IB-DNQ treatment in a mouse model with orthotopic xenograft. In a first proof of principle experiment, I injected the HSJD-DIPG-007 cell line into the pons area of 16 mice. I observed that IB-DNQ treatment significantly increased the survival of mice compared to the control group (**Fig. 5g**).

Overall, these data suggest that DIPGs carrying mutant truncated forms of PPM1D with associated overexpression of the stress related protein NQO1 are very sensitive to the anti-growth effect induced by the bioactivatable drug IB-DNQ, a molecular weakness that it is worth to be explored in further studies aimed to improve the survival of patients with currently dismal prognosis.

## Discussion

Herein I have observed NQO1 protein overexpression in DIPG patient-derived cell lines and primary tissues with PPM1D truncation. The best marker that discriminated the NQO1 overexpressing DIPG cells was the PPM1D mutation. By different cellular models, I deciphered that MDM2 S395 phosphorylation targeted by PPM1D, led to NQO1 accumulation. However, further studies would be needed to know more precisely how these proteins interact all together in the interplay between p53, PPM1D, MDM2 and NQO1 protein accumulation. The fact that MDM2 has been shown to promote translation of few mRNAs give rise to the hypothesis that it could also be the case for NQO1 mRNA in this specific context. The found PPM1D-mediated overexpression of NQO1 prompted us to use the NQO1-bioactivatable compound IB-DNQ in different DIPG models. The drug demonstrated a great activity against DIPG cell lines that was indeed dependent on NQO1 overexpression induced by the PPM1D truncated mutation. This finding suggests that DIPG patients carrying a PPM1D mutation could directly benefit from the use of this compound, or similar ones, a candidate tailored treatment that will require further extensive validation. However, for a tumour with such an extremely poor overall survival, the described data is encouraging for the design of more complete preclinical studies that could pave the way for an eventual clinical application.

The use of bioactivatable NQO1 compounds could also fit with the proposed treatment options for patients with DIPGs, since various have been described as modulators of NQO1 expression [12,30,37,41], like radiotherapy that is the standard of care in DIPG [6]. This is also the case for Gemcitabine, Doxorubicin, Cisplatin and Vorinostat [7,37,41]. Targeting NQO1 would not impair current management but could permit combinations of drugs that have been described to increase the expression of NQO1 potentiating the efficacy of IB-DNQ treatment. From the clinical standpoint of patient management and monitoring, the minimally invasive biopsy of DIPG could provide valuable information about driver mutations or protein overexpression that can lead to specific treatment. This would be the case of patients with PPM1D truncations linked to NQO1 upregulation, a group that could benefit from the administration of IB-DNQ or similar drugs. Importantly, the low toxicity of the compound described so far in animal models, combined with the extremely high need for new therapeutic options in DIPG, make research endeavours in this subject a promising area to address the unmet medical need of improving survival for this fatal childhood brain tumor.

## Authoŕs Contributions

**Conception and design:** M. Janin.

**Development of methodology:** M. Janin.

**Acquisition of data (provided animals, acquired, and managed patients’ data, provided facilities, etc.):** M. Janin.

**Analysis and interpretation of data (e.g., statistical analysis, biostatistics, computational analysis):** M. Janin.

**Writing, review and/or revision of the manuscript:** M. Janin.

**Administrative, technical, or material support (i.e., reporting or organizing data, constructing databases):** M. Janin.

**Study supervision:** M. Janin.

## Acknowledgments

I would like to thank Dr. Andres Morales La Madrid, Dr. Angel Montero Carcaboso and Dr. Jaume Mora from the Pediatric Cancer Center Barcelona (PCCB) at Hospital Sant Joan de Déu, Barcelona for their significant support. I think Dr. Michelle Monje from Stanford Medicine for providing SU-DIPG-VI cell line and SU-DIPG-35 frozen tissue and matched normal cortex. I warmly thank Dr. Pratiti Bandopadhayay, Dr. Eric Morin, and Dr. Timothy Phoenix from the Boston Dana-Farber Cancer Institute for providing data regarding IUE mouse models. I acknowledge Dr. Chris Jones from the Institute of Cancer Research, London, for sharing somatic mutations data information regarding DIPG cell lines. I thank Dr. Myung Ryul Lee and Dr. Paul J. Hergenrother for providing IB-DNQ compound and UZH’s DMG Research Center, Universitäts-Kinderspital Zürich for providing tumor samples from patients, especially Dr. Sulayman Mourabit, Dr. Javad Nazarian, and Dr. Denise Morinigo from the Children’s National Hospital, Washington D.C. I also thank CERCA Program / Generalitat de Catalunya for their institutional support.

## SUPPLEMENTARY INFORMATION

Supplementary Information include two Figures and one Key Resources Table.

Document S1. Key Resources Table.

**Figure S1.**
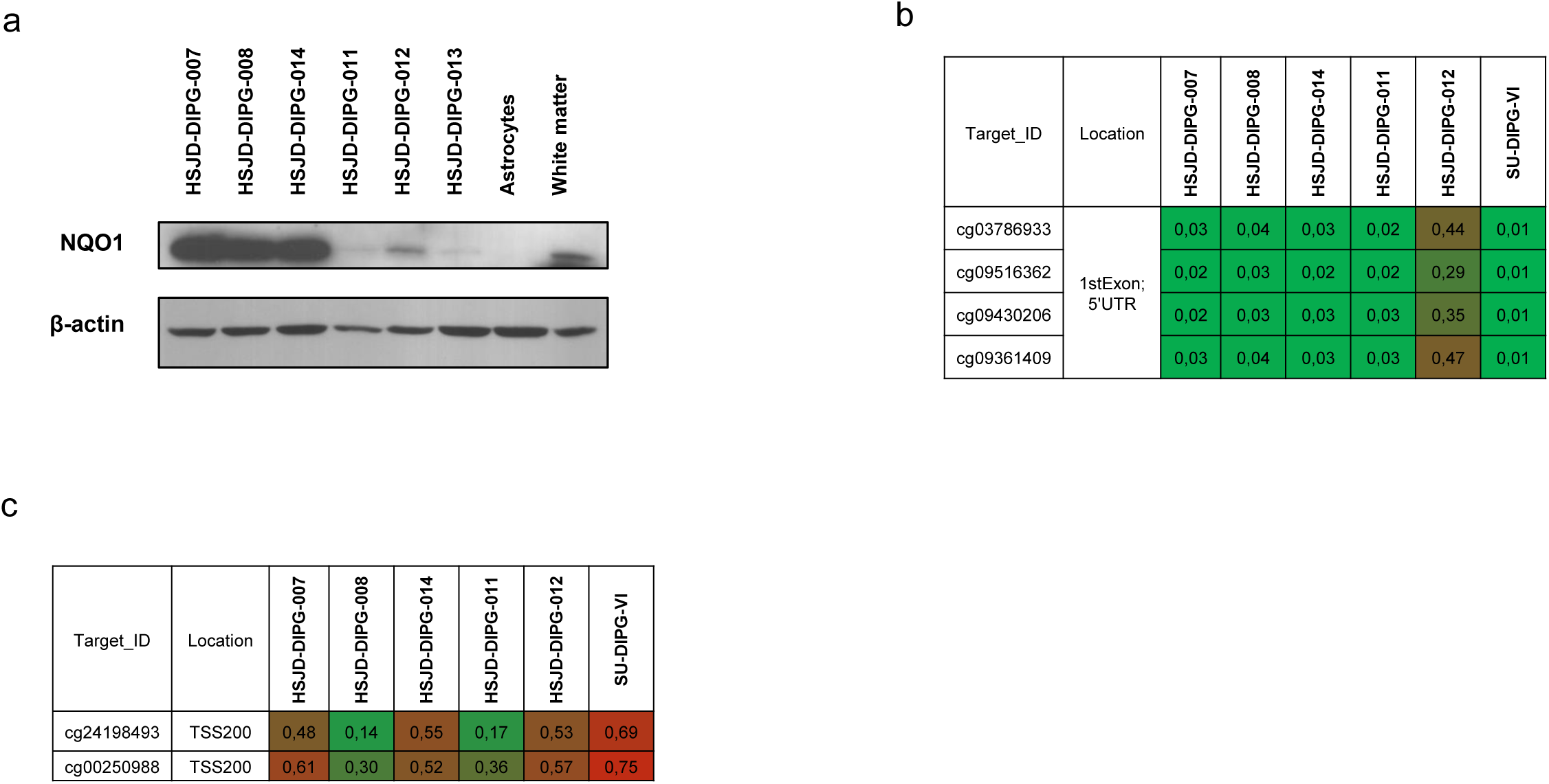
Related to Figure 1. (**a**) Western blot of NQO1 in 6 DIPG cell lines (containing HSJD-DIPG-013 but without SU-DIPG-VI) and normal human astrocytes or normal human white matter as controls. (**b**) DNA methylation profile of the CpG island promoter for the *NQO1* gene analysed by the EPIC DNA methylation microarray. Single CpG absolute methylation levels (0 – 1) are shown. Green, unmethylated; red, methylated. (**c**) DNA methylation profile of the CpG island promoter for the *NSUN5* gene.

**Figure S2.**
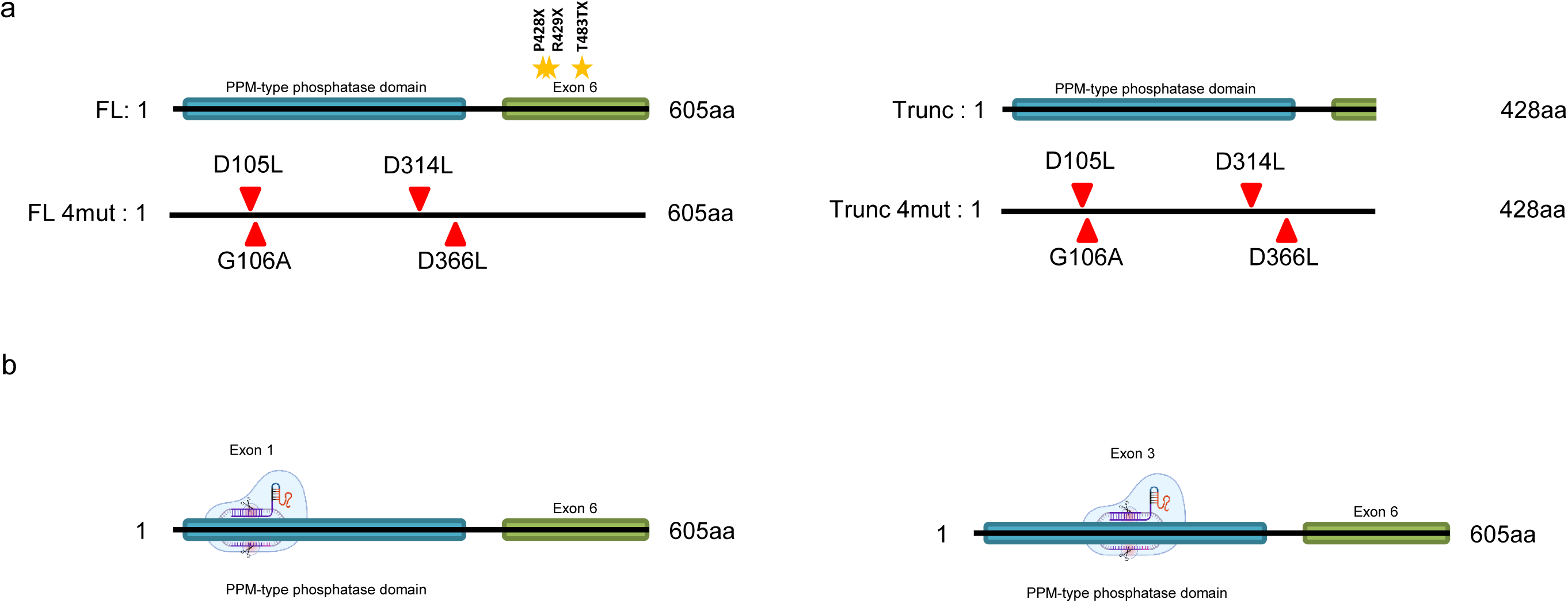
Related to Figure 3. (**a**) Mutational constructs in PPM1D transcripts for the full length form (FL ; left) or truncated (Trunc; right) that are wild type (up) or mutated (4mut; down). (**b**) Construction of the CRISPR-Cas9 PPM1D model in HSJD-DIPG-007 at exon 1 (left) or exon 3 (right).

**Key Resources Table.**
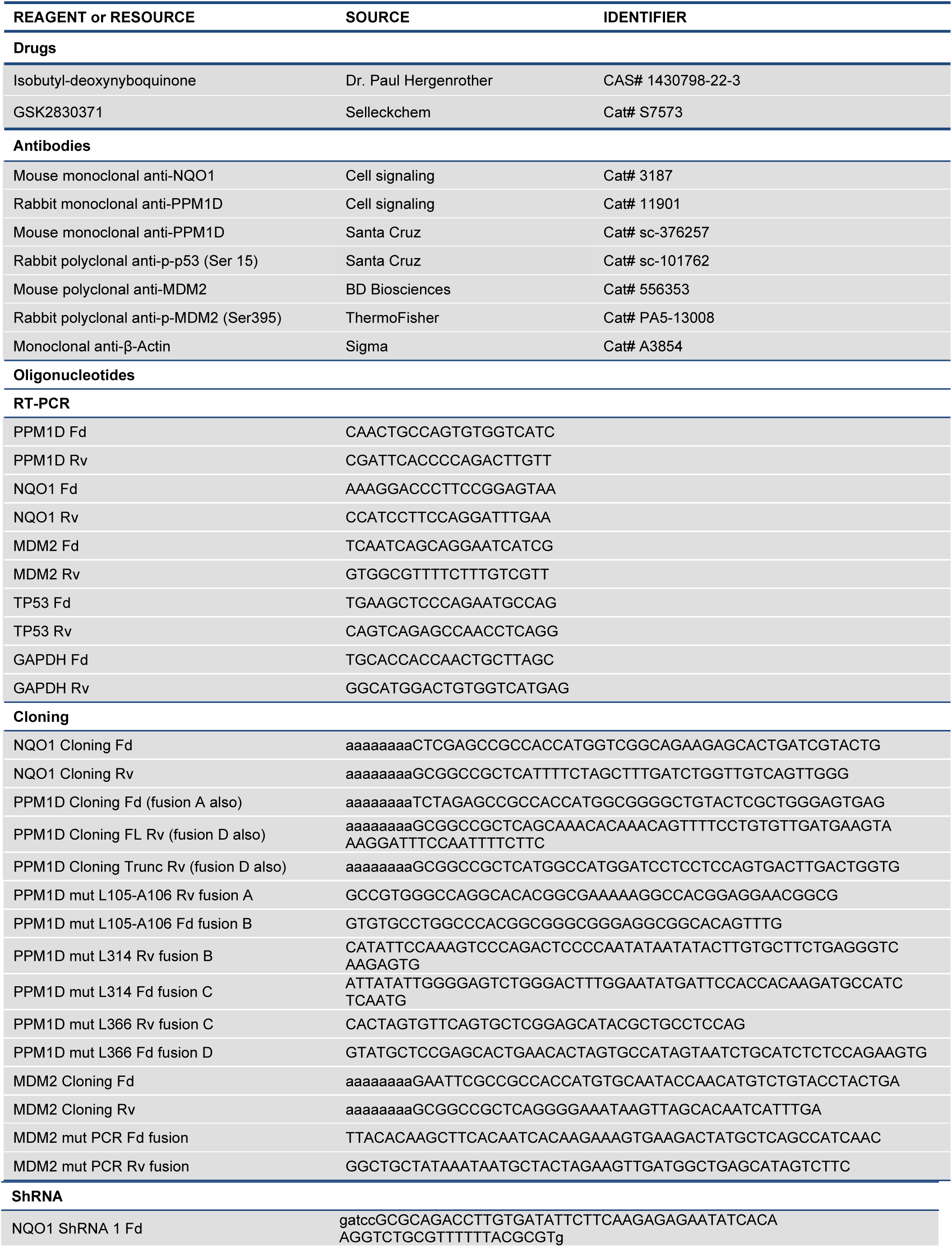
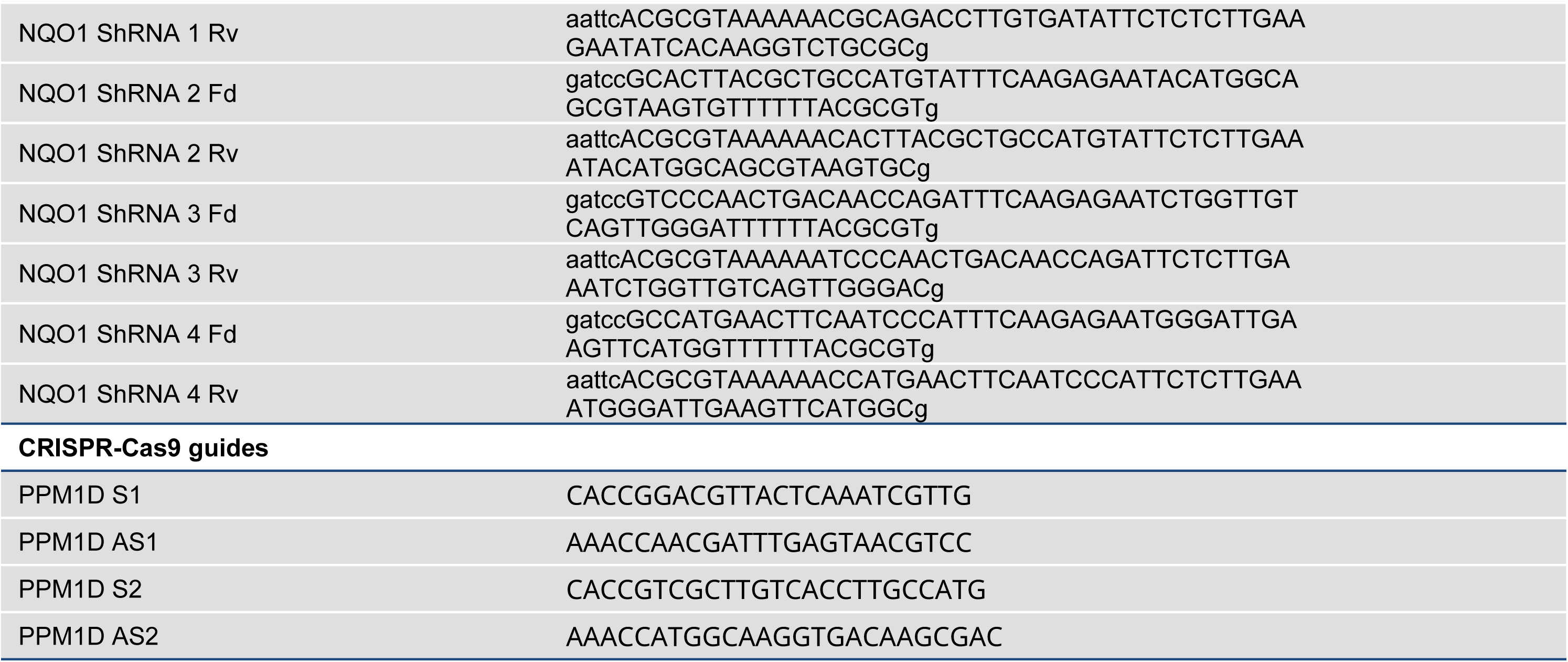

